# Mendel, a foundation model of human genetic variation, prioritizes regulatory variants and improves gene expression prediction

**DOI:** 10.64898/2026.09.28.755183

**Authors:** Arham Salman, Haonan Feng, Peng Wei, Ryan Sun, Lang Wu, Wei Pan, Chong Wu

## Abstract

Human genome-wide association studies have identified hundreds of thousands of variant trait associations, but interpreting their mechanistic consequences at allele resolution remains a central bottleneck. DNA foundation models pretrained on genomic sequence corpora learn features predictive of regulatory activity, yet their training objectives are dominated by the conserved background common to all individuals, rather than the sparse variant sites at which human genetic variation is concentrated. Here we introduce Mendel, a DNA foundation model trained from scratch on human whole genome sequences using a variant-centric objective that concentrates supervision on allelic events within reference aligned sequence context. On held-out individuals, Mendel attains 0.984 micro-averaged SNV allele-prediction accuracy, compared with 0.311 for Evo 2. Mendel residual difficulty is selectively enriched at cis-eQTL variants, prioritizes fine-mapped causal variants (AUROC 0.731), where Evo 2 residuals performed near chance, and when converted into a cis-expression prior for modelling, improves prediction relative to an unweighted baseline in all 50 GTEx tissues.

## Introduction

The discovery of genetic associations increasingly outpaces their biological interpretation. Genome-wide association studies and large-scale sequencing efforts have identified hundreds of thousands of variant-trait associations^1^, yet the consequences of most signals remain unresolved. This challenge is particularly acute because many implicated variants are noncoding, and their effects are often mediated through gene regulation^2^’^3^ and depend on biological context. Models that relate allelic differences to regulatory effects and molecular phenotypes could therefore help interpret these signals. Because genomic sequence contains part of the information needed to model these relationships, DNA foundation models (DNA-FMs) offer a scalable means of learning representations from sequence.

Through pretraining on large genomic sequence corpora^4–9^ or sequence-linked functional data^10–16^, these models capture aspects of regulatory grammar that can support downstream tasks like variant effect prediction. However, their objectives are typically defined densely across genomic positions, whether through next nucleotide prediction or functional genomic targets, so the training signal largely reflects the conserved sequence background shared among individuals. Human genetic variation, by contrast, is sparse relative to this sequence background and confined to the comparatively small subset of positions at which individuals differ. Variant effects in current DNA-FMs are consequently evaluated mainly after training, by contrasting predictions or embeddings for reference and alternate alleles, rather than serving as explicit targets during representation learning. Existing DNA-FMs are therefore powerful for understanding genomic sequence, but are not designed primarily to learn the structure of allelic variation across personal genomes^17,18^. This mismatch leaves a central gap for human genetics. The field still lacks a foundation model in which observed allelic variation itself is the primary object of learning and surrounding sequence provides context. Aligning the objective with the sparse, individual-level structure of human variation would also enable a direct test of whether the remaining prediction errors carry information relevant to variant function.

Here we introduce Mendel, a DNA-FM trained from scratch on human whole genome sequencing (WGS) data using a variant-centric objective. We named the new DNA-FM Mendel after Gregor Mendel, whose work laid the foundations of modern genetics. Mendel represents diploid genotypes as explicit events within reference-aligned sequence and concentrates the training loss on their allelic content (**Figure 1a**). We evaluate chromosome specific models at variant loci, asking whether learned population level patterns transfer to held out individuals. Mendel predicts the allelic content of held out individuals’ diploid genotypes at these cohort-defined loci more accurately than Evo 2. We then examine the structure and utility of the remaining prediction error. Residual difficulty is elevated at molecular quantitative trait loci (QTLs), supports discrimination of high posterior inclusion probability (PIP) cis-expression QTLs (cis-eQTLs) from low-PIP controls matched on minor allele frequency (MAF) and transcription start site (TSS) distance and, when used to weight variants, improves cis-expression prediction relative to an unweighted baseline in all 50 Genotype-Tissue Expression (GTEx) tissues. Together, these results position variant-centric modelling as a complement to general-purpose DNA sequence models and show that population-derived prediction residuals can provide a useful functional signal.

**Figure 1.**
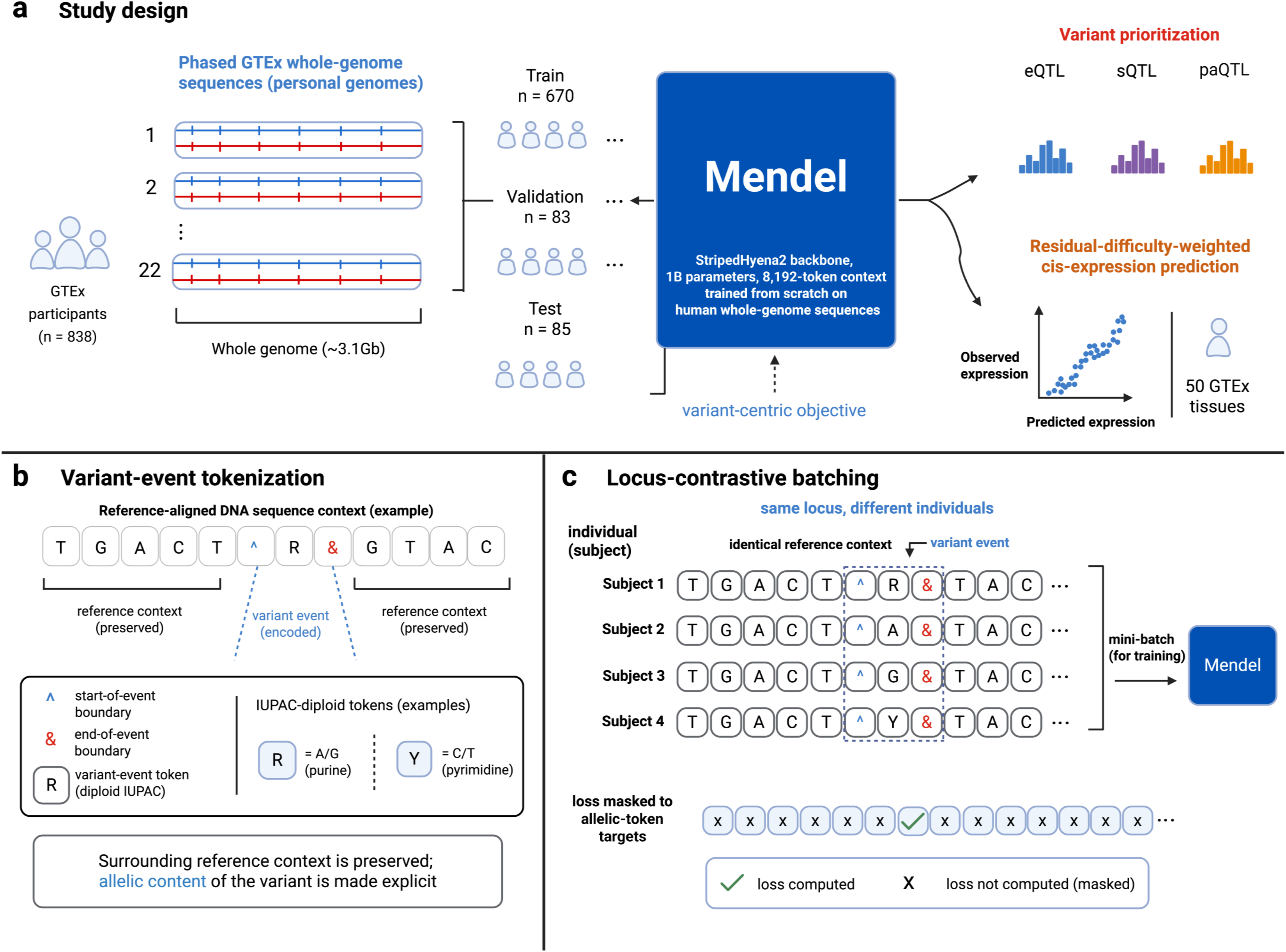
| A variant-centric foundation model of human genetic variation. (a) Study design. Mendel is trained from scratch on phased GTEx whole-genome sequences and evaluated on held-out individuals. Downstream applications span variant prioritization (QTLs) and residual-difficulty-weighted cis-expression prediction. (b) Variant-event tokenization. Each variant site is encoded as a compact IUPAC diploid token flanked by ^ and & boundary tokens, embedded within reference-aligned sequence context. (c) Locus-contrastive batching. Minibatches pair windows from the same local genomic region across different individuals, exposing Mendel to many allelic realizations of each context per gradient step. The training loss is masked to positions within ^…& boundaries.

## Results

### A variant-centric foundation model of human genetic variation

Mendel uses the StripedHyena2^19^ architecture and base nucleotide tokenizer of Evo 2 (1 billion parameters; 8,192 bp context)^6^, but is trained from scratch with a representation and objective designed around human genetic variation (**Methods**). Two design choices are central. First, Mendel encodes variants as discrete events within reference-aligned sequence (**Figure 1b**). At each single nucleotide variant (SNV) site, a single International Union of Pure and Applied Chemistry (IUPAC) ambiguity code encodes the diploid genotype (for example, R for A/G, Y for C/T)^20^. Two existing vocabulary tokens, ^ and &, delimit the start and end of each variant event. This representation preserves the local reference context while making allelic content explicit, captures both alleles without doubling sequence length and accommodates unphased genotypes because each IUPAC code denotes an unordered allele pair. Variable length content between the boundary tokens also permits representation of multi nucleotide variants and short indels. Second, Mendel uses locus-contrastive sampling. Each synchronized batch draws donor-specific windows from a common locus group, repeatedly exposing the model to different allelic realizations of the same local genomic region (**Figure 1c**). Like Evo 2, Mendel is trained autoregressively to predict the next token. In Mendel, however, cross-entropy loss is computed only for allelic tokens within variant-event boundaries; non-target tokens contribute no direct loss but provide preceding sequence context for each allelic prediction (**Methods**). Together, these choices focus optimization on predicting allelic content at ascertained variant loci from preceding sequence context.

We trained Mendel from scratch for each chromosome using WGS data from GTEx^21^ (**Methods**). Before training, the 838 individuals were randomly partitioned into disjoint training (n = 670), validation (n = 83), and test (n = 85) cohorts. Unless otherwise noted, every metric reported in this paper was computed exclusively on the held-out test cohort. We also evaluated the released 1 billion-parameter Evo 2 checkpoint using its published pretrained weights^6^ as an external reference for a current general-purpose DNA-FM.

### Mendel predicts genotype at variant sites in held-out individuals

Mendel is designed to learn a population-level representation of allelic variation at the variant sites on each chromosome. The evaluation therefore tests generalization to new individuals at sites represented during training, rather than prediction at unseen genomic positions. At each evaluated variant event, the task is to recover the held-out individual’s diploid genotype token from the preceding subject-specific genome sequence.

Across randomly selected eligible variant events on each chromosome, Mendel achieved a micro-averaged allele-prediction accuracy of 0.984, compared with 0.311 for Evo 2 (Δ = +0.673; **Figure 2a**). Evo 2, pretrained on 1 trillion nucleotide tokens from the phylogenetically diverse OpenGenome2 corpus, was evaluated through the same per-subject, locus-anchored pipeline and achieved substantially lower accuracy than Mendel. Thus, Mendel’s superior performance relative to Evo 2 highlights the value of explicit allele-focused training with variant-event representations for cross-individual genotype prediction.

**Figure 2.**
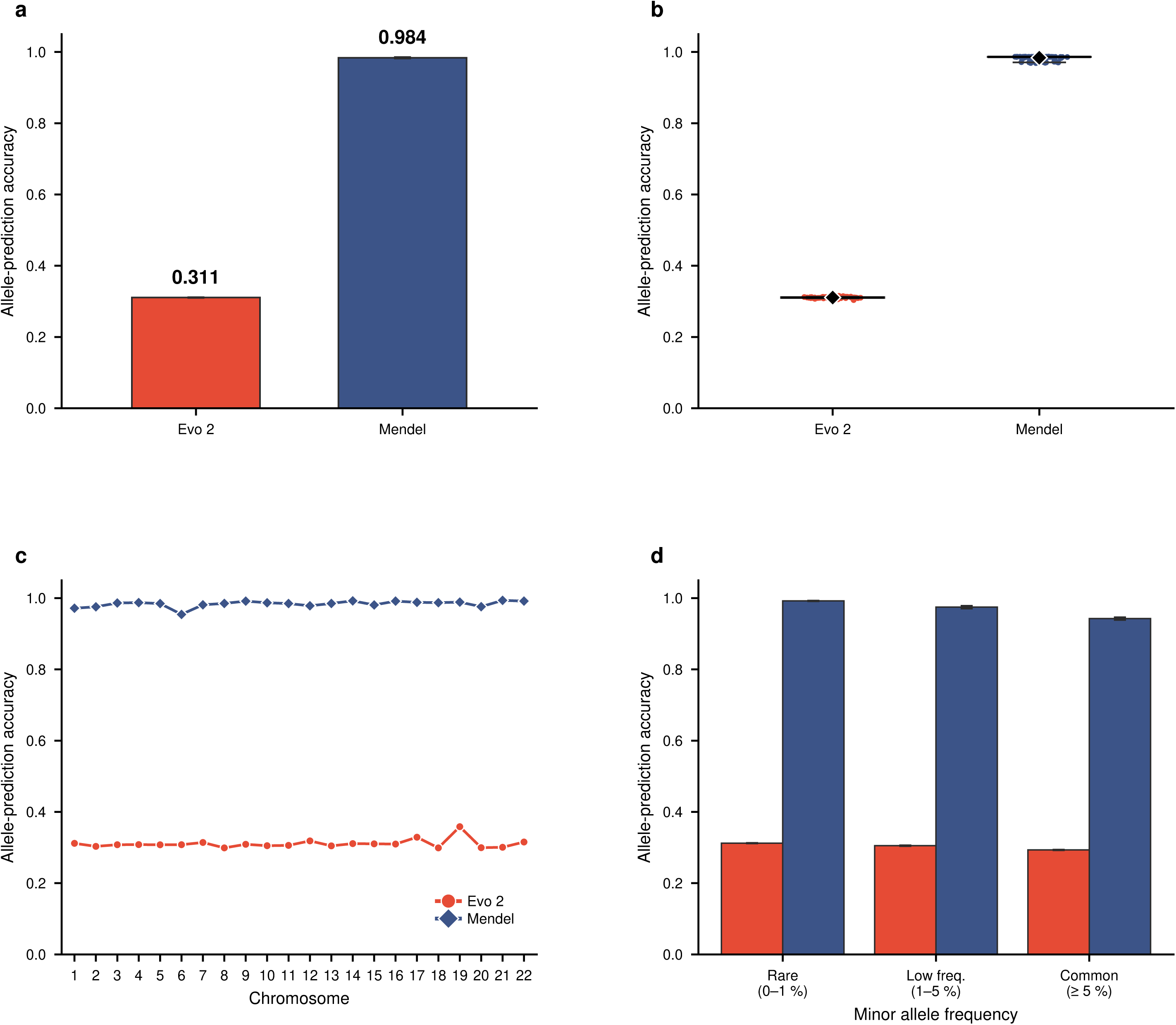
| Allele prediction in held-out individuals. (a) Micro-averaged allele-prediction accuracy for Evo 2 and Mendel, with 95% subject-bootstrap confidence intervals; labels give point estimates of 0.311 and 0.984. The models used the same donors and eligibility rules but independently sampled retained records. (b) Per-subject accuracy across the 85 held-out individuals; diamonds denote means and horizontal lines denote medians. (c) Allele-prediction accuracy by chromosome; the median Mendel - Evo 2 difference is +0.678. (d) Accuracy within rare (0-1%), low-frequency (1-5%) and common (≥ 5%) MAF bins, with 95% confidence intervals.

Mendel’s advantage over Evo 2 was consistent across individuals and chromosomes. Per-subject allele-prediction accuracy clustered near 0.31 for Evo 2 and near 0.98 for Mendel, and Mendel exceeded Evo 2 for each of the 85 held-out individuals (**Figure 2b**). Mendel also exceeded Evo 2 on every chromosome, with per-chromosome accuracy ranging from 0.95 to 0.99 for Mendel and from 0.30 to 0.36 for Evo 2 (**Figure 2c).** This chromosome-wide advantage shows that the aggregate result was not driven by a small subset of chromosomes.

Because micro-averaged accuracy can be sensitive to the allele-frequency spectrum and site composition of the evaluation set, we examined robustness under an alternative aggregation. Macro-averaging by variant, which gives each site equal weight, preserved Mendel’s advantage over Evo 2 (0.983 versus 0.311; **Extended Data Figure 1**). Mendel remained more accurate than Evo 2 within each MAF bin. Allele-prediction accuracies for Mendel and Evo 2, respectively, were 0.99 and 0.31 for rare variants (MAF < 1%), 0.97 and 0.31 for low frequency variants (1% ≤ MAF < 5%), and 0.94 and 0.29 for common variants (MAF ≥ 5%; **Figure 2d**). Genotype-stratified analysis showed that Mendel’s heterozygote allele-overlap score increased with MAF, from approximately 0.50 for rare variants to 0.62 for low-frequency variants and 0.92 for common variants. For Evo 2, heterozygote allele-overlap scores were approximately 0.29 across the three bins, reflecting single-allele recovery at a subset of sites **(Extended Data Figure 2).** To illustrate the contribution of genotype-class imbalance, we evaluated a descriptive, model-free predictor that returned two copies of the major allele at each site, with the major allele estimated from the held-out records. By construction, its allele-prediction accuracy was 1.0 for homozygous-major genotypes, 0.5 for heterozygous genotypes and 0 for homozygous-minor genotypes across MAF bins (**Extended Data Figure 3a**). The high aggregate accuracy in the rare-variant bin therefore does not imply strong recovery of rare non major genotypes, because homozygous major targets dominate that bin.

### Residual prediction difficulty is enriched at molecular QTLs

Although Mendel achieves high aggregate allele prediction accuracy, its remaining prediction errors were concentrated at a distinct subset of variants. We quantified these errors using hard miscall. For each subject-variant pair, hard miscall is the number of allelic mismatches between the most probable and observed diploid genotypes, with genotypes treated as unordered allele pairs; the score therefore ranges from 0 to 2 (**Methods**). Averaging this score across held out individuals defines the residual difficulty of each variant. Across the sampled evaluation variants, residual difficulty was strongly right skewed. Most variants had little or no residual error, whereas a smaller subset formed a long tail of more difficult sites (**Figure 3a**; **Extended Data Figure 4a**).

**Figure 3.**
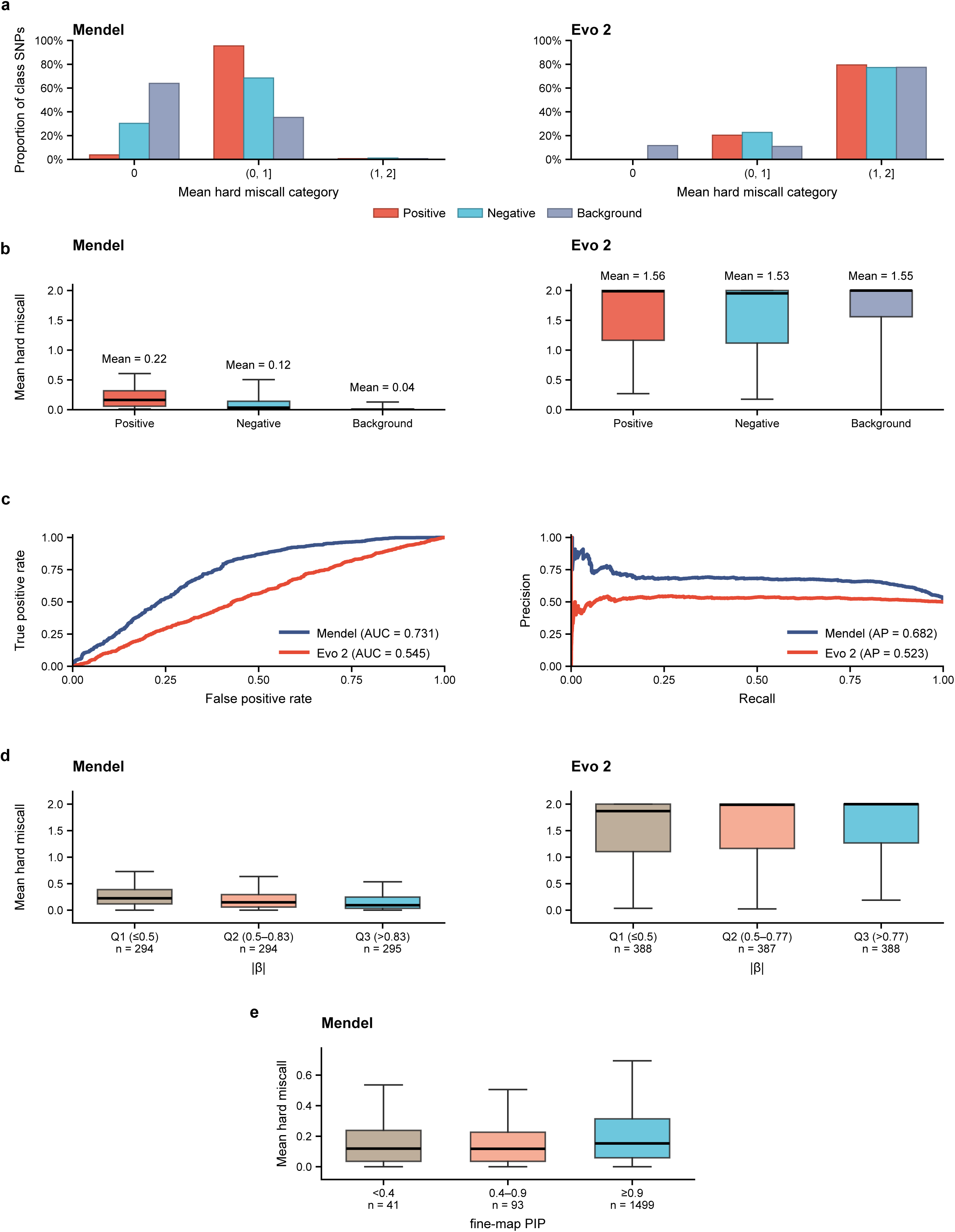
| Residual allele-prediction difficulty highlights functional variants and supports prioritization. (a) Proportions of cis-eQTLs (Positive), MAF- and TSS-distance-matched low-PIP controls (Negative) and random background variants in mean-hard-miscall categories 0, (0, 1] and (1, 2], shown separately for Mendel and Evo 2. Matching was performed separately for each model. (b) Mean per-variant hard miscall for the same three classes under each model; boxes show interquartile ranges and whiskers show the 5th-95th percentiles. (c) Receiver operating characteristic (ROC) and precision-recall curves from shuffled, stratified fivefold variant-level cross-validation distinguishing high-PIP cis-eQTLs from matched low-PIP controls (Mendel AUROC = 0.731, AP = 0.682; Evo 2 AUROC = 0.545, AP = 0.523). Model estimates derive from separately matched sets. (d) Residual difficulty versus absolute eQTL effect size in the matched positive subsets (Mendel: ρ = -0.308, *P = 8.5 ×* 10^-21^, n = 883; Evo 2: ρ = 0.083, *P* = 0.005, n = 1,163). (e) Mendel residual difficulty versus fine-mapping PIP in the unfiltered positive set (ρ = 0.156, *P* = 2.5 × 10^-10^, n = 1,633).

We first asked whether cis-eQTLs were enriched among variants with high residual difficulty. We used cis-eQTLs and low-PIP controls from a predefined benchmark released with the Borzoi study^15^. Positive variants had fine-mapping PIP of at least 0.9, and the corresponding released low-PIP variants formed the candidate control pool. Because residual difficulty was strongly associated with MAF, we matched positives and controls on MAF and distance to the nearest TSS, separately for Mendel and Evo 2 (**Methods**; **Extended Data Figure 5**); background variants were sampled independently. Under Mendel, residual difficulty followed a clear gradient: cis-eQTLs had the highest difficulty (median 0.17, mean 0.22), followed by matched controls (median 0.04, mean 0.12) and random background variants (median 0.00, mean 0.04; two-sided Mann-Whitney U tests: cis-eQTLs versus controls, *P = 2.8 ×* 10^-61^; controls versus background, *P* = 6.3 × 10^-120^; **Figure 3b**). In comparison, under Evo 2, hard-miscall scores were concentrated near their upper limit in all three variant groups (medians approximately 2.0; 90.0% of cis-eQTLs, 87.9% of controls and 79.7% of background variants scored above 0.5), and the cis-eQTL and control distributions were similar (**Figure 3b**, right). Thus, Mendel residual difficulty was enriched at cis-eQTLs relative to both the MAF- and TSS-distance-matched controls and the randomly sampled background set, whereas Evo 2 did not show comparable separation. Residual difficulty was also elevated at splicing, polyadenylation and intronic-polyadenylation QTLs relative to their released, mechanism-specific Borzoi controls (**Extended Data Figure 6**).

Because residual difficulty could vary with allele frequency and genomic context, we assessed these relationships directly. Residual difficulty was positively associated with MAF (Spearman ρ = 0.603, *P* = 3.6 × 10^-187^). Associations with distance to the nearest TSS (ρ = -0.262, *P* = 2.1 × 10^-31^) and local guanine-cytosine content (ρ = 0.238, *P* = 4.0 × 10^-26^) were smaller in magnitude (**Extended Data Figure 7**). Despite this MAF association, the cis-eQTL-control difference remained evident within MAF strata (median differences = 0.006, 0.012 and 0.188 for rare, low-frequency and common variants) and within TSS distance strata (median differences approximately 0.12 and 0.14 for near and intermediate variants; **Supplementary Table S1**). The difference was also preserved when residual difficulty was quantified using expected miscall, a probability weighted average of the distance between the observed genotype and each possible genotype call (Methods; **Extended Data Figure 8**).

We next asked whether this group level separation could support variant level cis-eQTL prioritization. Random forest^22^ classifiers were trained to distinguish high-PIP cis-eQTLs from matched low-PIP controls using residual-derived features, covariates comprising MAF, distance to the nearest TSS and chromosome indicators, or their union (Methods). In shuffled, stratified fivefold variant level cross-validation, Mendel residual features achieved an area under the receiver operating characteristic curve (AUROC) of 0.731 and average precision (AP) of 0.682, compared with an AUROC of 0.545 and AP of 0.523 for Evo 2 residual features (**Figure 3c**). As a sensitivity analysis, adding MAF, TSS distance and chromosome indicators did not materially improve performance over residual features alone **(Extended Data Figure 4b,c).**

Because the residual features are estimated across individuals, we examined prioritization performance as the number of contributing subjects varied (**Extended Data Figure 3b**). Using residual-only classifier, performance with eight subjects (mean AUROC approximately 0.74; mean AP approximately 0.69) was similar to that obtained with all 85 subjects (AUROC approximately 0.74; mean AP approximately 0.70), with little change at intermediate sample sizes. Evo 2 performance remained near an AUROC of 0.5 throughout. Mendel-derived prioritization was therefore stable across the subject counts examined.

We next examined whether Mendel’s residual difficulty tracks quantitative measures of regulatory effect and fine-mapping confidence (**Figure 3d,e**). Within the model-specific matched positive subsets used for **Figure 3d**, residual difficulty was negatively correlated with absolute eQTL effect size under Mendel (Spearman ρ = -0.308, *P* = 8.5 × 10^-21^, n = 883) but showed a weak positive correlation under Evo 2 (ρ = 0.083, *P* = 0.005, n = 1,163). In the unfiltered positive set for Mendel used for **Figure 3e**, residual difficulty was positively correlated with fine-mapping PIP (ρ = 0.156, *P* = 2.5 × 10^-10^, n = 1,633). The opposing signs of the Mendel correlations (difficulty decreasing with effect magnitude but increasing with fine-mapping PIP) suggest that residual difficulty captures a dimension of regulatory architecture aligned with fine-mapping evidence rather than simply tracking effect magnitude.

### Residual-difficulty priors improve cis gene-expression prediction

We next asked whether Mendel’s residual difficulty could be repurposed as an annotation-free functional prior for predicting gene expression from genotype data. Within each outer training fold, the mean hard-miscall score for each cis-window variant was converted into a gene-specific, rank-based elastic-net penalty factor, such that variants with greater residual difficulty underwent less shrinkage (**Methods**)^23^’^24^. Expression measurements were not used to construct these priors.

For this downstream analysis, donor sets were determined separately for each gene - tissue pair using all subjects with aligned genotype, residual-difficulty and tissue-specific expression data, rather than being restricted to the 85-donor test cohort. We evaluated 110 candidate genes randomly selected across the chromosomes (five per chromosome); 105 met the data-availability criteria in at least one tissue, yielding 4,896 evaluable gene-tissue pairs across 50 GTEx tissues. Within each tissue, the primary baseline comparison was restricted to the union of genes with prediction significant after false discovery rate (FDR) correction under either the baseline or Mendel-weighted model. Mendel weighting increased mean squared cross-validated Pearson correlation in all 50 tissues (**Figure 4a**), corresponding to a median relative increase of 3.93% (**Figure 4b**), and yielded more FDR-significant genes than the baseline model in 45 tissues (**Figure 4c**).

**Figure 4.**
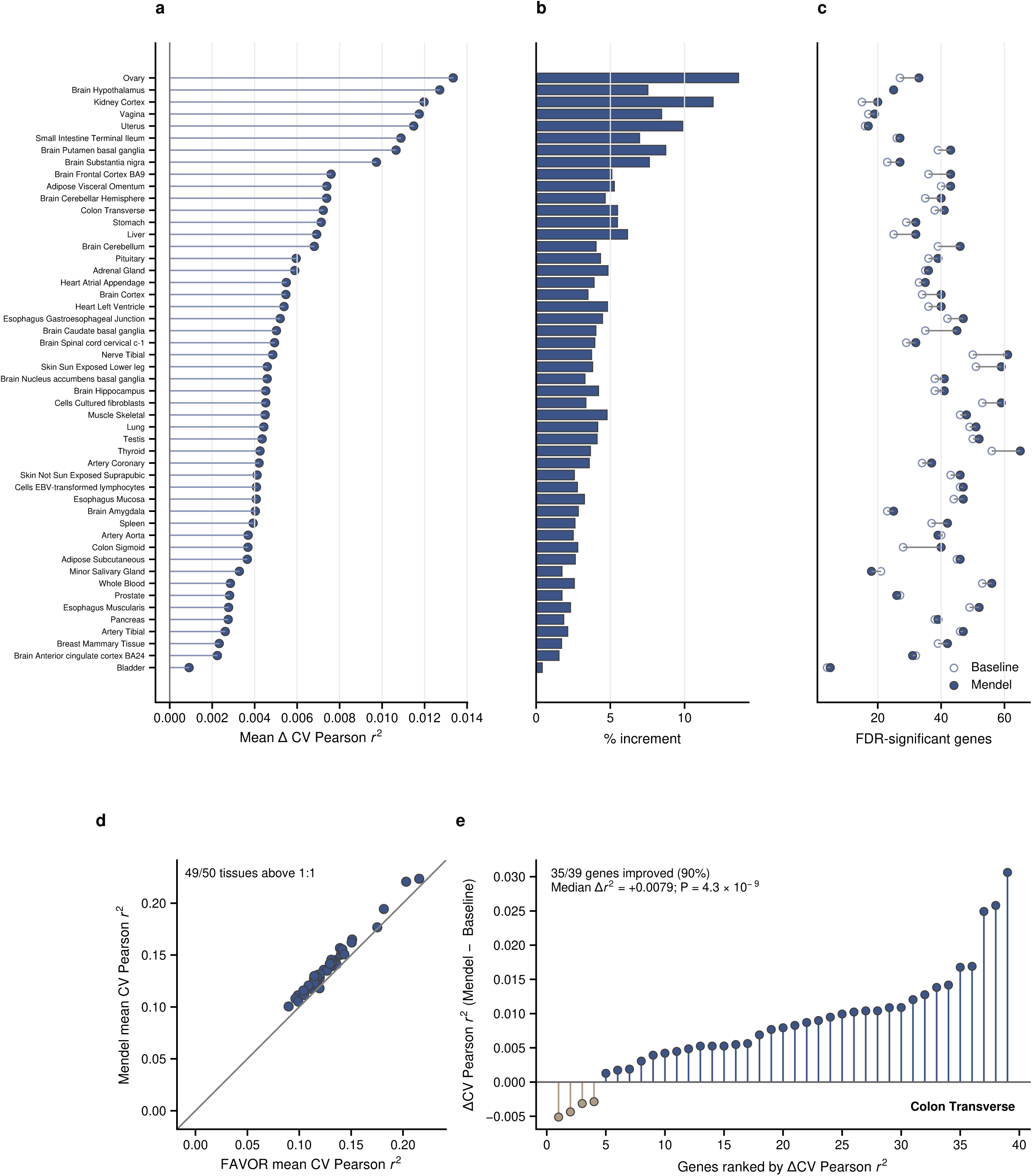
| Residual-difficulty priors improve cis gene-expression prediction. (a) Tissue-specific mean gain in squared cross-validated Pearson correlation for Mendel-weighted relative to baseline elastic-net models, evaluated among genes with FDR-significant prediction under either model. (b) Corresponding percentage improvement relative to baseline, with tissues shown in the same order as in a. (c) Numbers of FDR-significant genes (q ≤ 0.05) under the baseline and Mendel-weighted models in each tissue. (d) Tissue-level mean squared cross-validated Pearson correlation under Mendel residual-difficulty weighting versus CADD-Phred weighting obtained through FAVOR, evaluated among genes significant under either strategy; the dashed line denotes equality. (e) Gene-level performance in Colon Transverse among genes with mean cross-validated Pearson r > 0.10 under either the baseline or Mendel-weighted model; 35 of 39 genes had higher squared correlation under Mendel weighting.

To benchmark the informativeness of Mendel’s residual against a widely used annotation-based prior, we next compared Mendel residual-difficulty weighting with a prior based on PHRED-scaled Combined Annotation Dependent Depletion (CADD), a score that integrates diverse genomic annotations to rank variants by predicted deleteriousness. CADD-Phred annotations were obtained through Functional Annotation of Variants Online Resource (FAVOR)^25^. Among genes with FDR-significant prediction under at least one strategy, mean squared cross-validated Pearson correlation was higher with Mendel weighting in 49 of 50 tissues (**Figure 4d**). Colon Transverse, selected for the gene-level view using the procedure described in Methods, contained 39 genes exceeding a mean cross-validated Pearson correlation of 0.10 under either the baseline or Mendel-weighted model. Mendel weighting improved squared correlation for 35 of these genes (median Δr^2^ = 0.0079; one-sided paired Wilcoxon signed-rank *P* = 4.3 × 10^-9^; **Figure 4e**). Thus, across the evaluated gene-tissue sets, Mendel residual-difficulty weighting improved cis-expression prediction and provided a stronger functional prior than CADD-Phred weighting.

## Discussion

Mendel recasts DNA-FMs around human genetic variation itself, rather than treating variation as a downstream perturbation of a reference centric sequence model. By representing each variant as an explicit event within conserved sequence context and concentrating supervision on allelic content, Mendel makes allele level prediction a primary capability rather than an emergent byproduct. In held out individuals, Mendel achieved substantially higher aggregate allele-prediction accuracy than Evo 2. More importantly, its remaining prediction errors were functionally structured: residual difficulty was enriched across multiple classes of molecular QTLs and supported prioritization of fine-mapped regulatory variants.

Because the training and test individuals share population structure, Mendel could perform well by learning the common allele at each site and using nearby variants as proxies through linkage disequilibrium (LD). Three downstream analyses demonstrate that Mendel’s residual difficulty captures functionally relevant variation beyond aggregate genotype-prediction accuracy. First, residual difficulty was elevated at cis-eQTLs relative to MAF and TSS distance-matched controls and at other molecular QTLs relative to their released, mechanism-specific Borzoi controls. Second, Mendel residual difficulty correlated negatively with eQTL effect size but positively with fine-mapping PIP. Third, Mendel-based weighting yielded higher mean squared cross-validated Pearson correlation than CADD-Phred weighting in 49 of 50 tissues. Together, these findings support the functional relevance of Mendel’s residual difficulty.

Mendel is best understood as complementary to, rather than a replacement for, existing DNA-FMs. Evo 2^6^ learns position specific sequence representations, whereas AlphaGenome^16^, Enformer^13^, and Borzoi^15^ predict regulatory activity across long genomic contexts. Mendel instead models sequence-to-variation relationships by learning how population allelic diversity depends on local genomic context. Sequence-to-function and sequence-to-variation models therefore capture complementary information about a variant: its predicted biological effect and fit within local patterns of population variation. Combining these signals, for example by integrating track-based effect predictions with Mendel’s residual-difficulty priors, could improve both mechanistic interpretation and downstream discovery. More broadly, Mendel’s results suggest that variant-centric objectives may be a generally productive strategy for aligning representation learning with the sparse, individual-level structure of human genetic variation, a direction likely to be increasingly relevant as biobank-scale sequencing cohorts continue to grow.

Five limitations define the next steps for extending Mendel. First, we trained a separate model for each chromosome to make locus-contrastive training computationally feasible. Each minibatch contains the same 8,192-bp locus from multiple individuals. Partitioning the genome by chromosome reduced the number of loci handled in each training run while preserving dense cross-individual sampling at every locus. Scaling Mendel to a single genome-wide model is a natural next step and would enable trans chromosomal inference, though at substantially higher compute cost. Second, Mendel was evaluated at variant loci ascertained in the full cohort, so generalization to previously unseen loci was not tested. Variant prioritization was evaluated using randomly assigned variant-level folds. Future locus or LD block-held-out evaluation will be important for assessing generalization across independent genomic regions. Third, Mendel’s current training cohort is GTEx, whose ancestry composition does not reflect global human diversity. Extending training and evaluation to cohorts such as All of Us^26^, the Pan-UK Biobank^27^, and the Million Veteran Program^28^ will be essential for understanding how variant-centric representations transfer across ancestries, and for supporting equitable downstream use. Fourth, because heterozygotes constitute only a small proportion of genotypes at rare sites, their poor recovery has little effect on micro-averaged accuracy, but limits applications focused on rare-variant carriers. Future models could address this limitation by assigning greater training weight to rare heterozygous genotypes. Fifth, Mendel is computationally intensive during training and inference, which limited systematic hyperparameter optimization. Broader tuning may therefore improve performance, while smaller architectures and optimized inference could reduce runtime and memory demands and improve scalability.

## METHODS

Methods are available in the **Methods** section.

*Note: One supplementary pdf file and one supplementary table file are available*.

## ACKNOWLEDGEMENTS

Research reported in this work was supported by National Cancer Institute under Award Numbers R01CA263494, U01CA293883, and P30CA016672, National Institute on Aging under Award Numbers U01AG073079, and Sabin Family Fellowship. The content is solely the responsibility of the authors and does not necessarily represent the official views of the National Institutes of Health.

## AUTHOR CONTRIBUTIONS

C.W. and W.P. designed the study. A.S. and C.W. developed Mendel. A.S. trained Mendel and analyzed the data. H.F. helped prepare data resources and analyze data and provided feedback on the study and platform design. W.P, R.S., and L.W provided valuable feedback to strengthen the design of the study. A.S. and C.W. wrote the manuscript with feedback from all authors.

## COMPETING INTEREST

The authors declare no competing interests.

## Methods

### Training data and sequence preparation

Genotypes were obtained from the SHAPEIT2-phased GTEx v8 whole-genome call set (GTEx_Analysis_2017-06-05_v8_WholeGenomeSeq_838Indiv_Analysis_Freeze), comprising 838 donors with variants represented relative to GRCh38. Although the source genotypes are phased, Mendel represents each diploid SNV genotype as a single unordered IUPAC symbol^20^ and therefore does not retain haplotype phase. A, C, G and T represent A/A, C/C, G/G and T/T, respectively; M, R, W, S, Y and K represent A/C, A/G, A/T, C/G, C/T and G/T.

A genomic locus was designated a variant if at least one of the 838 donors carried a non-reference allele at that locus. The resulting catalogue of variant loci was shared across all donors. At each locus in this catalogue, the donor’s genotype was enclosed by the ^ and & special tokens in that donor’s FASTA sequence. Thus, the same genomic loci were marked in the training, validation and test sequences, while the genotype encoded between the special tokens varied among donors.

Donors were randomly assigned to an 80/10/10 non-overlapping training, validation and test partitions using seed 42, yielding 670 training, 83 validation and 85 test donors, respectively. This partition was held fixed across chromosomes and all subsequent analyses, and all reported allele-prediction metrics were calculated using only the 85 test donors. Each donor-specific encoded chromosome was divided into overlapping 8,192-token windows with a stride of 4,096 tokens, resulting in 50% overlap between adjacent windows. Windows were retained if they contained at least one marked variant and equal numbers of opening (^) and closing (&) boundary tokens, and were subsequently grouped by genomic locus for locus-contrastive batching.

Donor ancestry composition is reported in Supplementary Table S4.

### Model architecture

Mendel adopts the StripedHyena2 backbone^19^ of Evo 2^6^ at 1 billion parameters, with a context length of 8,192 base pairs, unmodified otherwise. The configuration comprises 25 layers, hidden size 1,920, feed-forward size 5,120 and 15 attention heads, in Evo 2’s Hyena-SE / Hyena-MR / Hyena-LI / attention operator pattern. Initialization is random; no pretrained Evo 2 weights are used.

Mendel uses a byte-level tokenizer in which every input character is represented by its byte value. The ^ and & delimiters and the IUPAC symbols are already present in this vocabulary, so the variant-event grammar requires no vocabulary extension.

### Variant-event tokenization

At each variant site in a subject’s genome, we emit a single compact token encoding the diploid genotype under the IUPAC ambiguity code (for example, R for A/G, Y for C/T, S for C/G)^20^. This captures both alleles without doubling sequence length and, because each IUPAC symbol is defined by the unordered allele pair, is compatible with unphased genotype data. The variant token is flanked by the ^ (start) and & (end) boundary tokens, such that a single-nucleotide variant at position *x* is represented as the triplet ^ [IUPAC(x)] & embedded within the surrounding reference-aligned sequence. The encoding extends to multi-nucleotide variants and short indels by allowing variable-length content between the event boundaries.

### Locus-contrastive batching

Within training windows, the loss was applied only to genotype content in successfully paired ^…& events.

Each retained window was assigned to a locus group according to its start offset in the encoded chromosome. Windows with start offsets within the same 8,192-token interval were assigned to the same group. Because consecutive windows began 4,096 tokens apart, a locus group could contain overlapping windows with different start positions. A locus group therefore represented the same portion of the encoded chromosome rather than an identical genomic window.

Each synchronized sampler batch was drawn from one locus group. For every optimizer step, the sampler selected one eligible window from each of eight distinct donors. The runtime processed one Megatron microbatch per optimizer step, yielding an effective global batch size of eight windows; no gradient accumulation was used. Donors were unique within a synchronized batch but could appear in a subsequent batch if another eligible window remained. A locus group continued to produce batches only while at least eight donors had eligible windows available.

Locus order, donor order and candidate windows were shuffled using a NumPy random-number generator initialized with seed 42.

### Variant-only next token loss

Mendel was trained using autoregressive cross-entropy restricted to the genotype tokens within marked variant events. Each event was encoded as ^g&, where g denotes the donor’s IUPAC genotype token. Labels were shifted by one position for next-token prediction, and the loss mask was placed at the position of the opening boundary. Thus, for ^g&, the model predicted g from the preceding sequence ending in ^. Neither boundary token was itself a prediction target.

More generally, if the opening and closing boundaries occurred at token positions (s) and (e), the loss mask covered positions (s) through (e-2). Because each label was the following token, the corresponding targets were exactly the tokens between the two boundaries. Positions outside marked events contributed no direct loss but remained part of the causal input context.

All successfully paired events within a retained window contributed to the objective. Windows were indexed only when they contained at least one ^ token and equal numbers of ^ and & tokens. During mask construction, each opening boundary was paired with the next unused closing boundary to its right, and a pair contributed to the mask only when it enclosed at least one token. If no valid target remained, an all-zero loss mask was used rather than reverting to whole-sequence next-token prediction.

Within each per-GPU microbatch, cross-entropy was summed over the masked genotype targets and divided by the number of masked targets. The resulting microbatch losses were averaged during distributed optimization. Each genotype target therefore received equal weight, while windows containing more marked events contributed more loss terms. A microbatch with no masked targets contributed zero sequence-loss gradient, although the optimization step itself was not skipped.

### Per-chromosome training

A separate model was trained for chromosomes 1-22 using genotype-encoded sequences from the training donors and the cohort-defined catalogue of marked variant loci for that chromosome. Training used overlapping 8,192-token windows from the corresponding chromosome and the locus-contrastive batching procedure described above.

Each chromosome-specific run used four NVIDIA H100 GPUs, with tensor- and pipeline-parallel sizes of 1. Each optimizer step consisted of one synchronized Megatron microbatch, with two windows processed on each GPU. The effective global batch size was therefore eight windows, with no gradient accumulation. Training used bfloat16 mixed precision and a maximum budget of 200,000 optimizer steps. Training duration varied by chromosome. Runs for the smaller chromosomes (13-22) ended after approximately 50,000-81,000 optimizer steps. Among chromosomes 1-12, most runs ended after approximately 84,000-100,000 steps; chromosome 6 ended after approximately 50,000 steps, and chromosomes 10 and 12 continued to approximately 132,500 and 155,000 steps, respectively. The number of window presentations was eight times the number of completed optimizer steps and therefore varied across chromosomes, with a maximum of 1.6 million per run.

Optimization used Adam with *β1* = 0.9, β_2_ = 0.95 and *ε* = 10^-8^, weight decay of 0.001 and gradient-norm clipping at 5.0. The executed-run logs show a peak learning rate of 10^-4^. The scheduler used 100 warm-up steps to reach the peak rate, followed by cosine decay toward a minimum learning rate of 10^-5^(2×10^-5^ for chromosomes 13, 15, 17, 19 and 20), with a 500 step terminal hold at the minimum rate.

Validation loss was evaluated every 1,250 optimizer steps and used to monitor convergence. Checkpointing retained the five checkpoints with the lowest validation loss as well as the most recent checkpoint. The checkpoint used for each downstream analysis, together with its training step and full run configuration, is reported in **Supplementary Table S2**. Training and validation loss trajectories for the chromosome-specific models, together with the selected checkpoints, are shown in **Extended Data Figure 9**.

### Baselines

Evo 2 was evaluated using its released 1-billion-parameter, 8,192-token pretrained checkpoint without additional training. For each scored single-token SNV event, the marked sequence ^g& was transformed to g, where g denotes the donor’s genotype token, and position identifiers were regenerated for the transformed input. Prediction used the logits at the sequence position immediately preceding g. The target position was supplied by the locus-anchored evaluation; Evo 2 was therefore evaluated on genotype prediction at known variant sites rather than being required to identify the sites themselves.

As a model-free frequency reference, we evaluated a predictor that returned two copies of the major allele at each site. Truth records were pooled from the saved held-out evaluation outputs and deduplicated by donor, chromosome and genomic position. The major allele was estimated from these retained records. The predictor received an allele-overlap score of 1 for a homozygous-major genotype, 0.5 for a heterozygous genotype and 0 for a homozygous-minor genotype. Because the major allele was estimated from the held-out records used in this analysis, this baseline was treated as a descriptive reference for genotype-class imbalance rather than as a prospective predictor.

### Genotype-prediction accuracy

The model was evaluated at single-token genotype events represented as ^X&, where (X) represents an IUPAC coded variant. Up to 10,000 events were evaluated per donor on each chromosome. Evaluation was performed on the 85 held-out donors using 8,192-token windows with a stride of 4,096 tokens. Only complete, single-token SNV events of the form ^g& were scored. For Mendel, the opening boundary was required to occur at least 4,096 tokens from the beginning of the marked input window. For Evo 2, the corresponding prediction position in the transformed input was required to occur at least 4,096 tokens from the beginning of that sequence.

For each donor and chromosome, candidate windows were shuffled independently for Mendel and Evo 2 and processed in the resulting order. Eligible events were retained in encounter order until a maximum of 10,000 events was reached. Consequently, the retained event sets were selected independently for the two models, and no shared donor-event manifest was used.

At each event, the task was to predict the observed diploid genotype token from the preceding donor-specific sequence. Mendel prediction used the logits at the opening boundary and was restricted to the ten diploid SNV symbols A, C, G, T, M, R, W, S, Y and K. Evo 2 prediction used the logits immediately preceding g in the transformed input and was restricted to A, C, G and T.

Genotype symbols were decoded as unordered allele pairs. A, C, G and T represented homozygous genotypes, whereas M, R, W, S, Y and K represented the corresponding heterozygous genotypes. Predictions were scored by symmetric diploid allele overlap. Exact genotype agreement received a score of 1, recovery of one allele received 0.5 and recovery of neither allele received 0. For Evo 2, each A/C/G/T prediction was interpreted as the corresponding homozygous genotype; at a heterozygous target, a match to either observed allele therefore received a score of 0.5. Micro-averaged allele-prediction accuracy was the mean score across all retained records for each model. Per-donor and per-chromosome accuracies were calculated as the corresponding means within each donor or chromosome. Site-macro accuracy was calculated by first averaging scores at each chromosome-position pair and then averaging across sites, giving each retained site equal weight.

MAF was calculated from the full 838-donor call set. Frequency analyses were restricted to biallelic SNVs, defined as sites with one reference nucleotide and one alternative nucleotide. Variants were classified as rare (MAF < 1%), low frequency (1% ≤ MAF < 5%) or common (MAF ≥ 5%). Sites with multiple alternative alleles, insertions or deletions, or substitutions spanning more than one nucleotide were excluded from MAF-stratified analyses. Within each MAF category, records were further stratified by the observed genotype as homozygous minor, heterozygous or homozygous major. Heterozygotes were identified directly from their IUPAC symbols. For homozygous records, major- and minor-allele identity at each site was inferred from allele counts in the retained Evo 2 truth records and then applied to both models.

### Residual-difficulty scores

For the residual-difficulty analyses, each variant was evaluated once per donor using a single 8,192-token window centered on the variant. This benchmark was conducted separately from the sliding-window evaluation used to estimate genotype-prediction accuracy. For donor (s) and variant (v), let *(**p_sv_(g)**)* denote the constrained probability assigned to diploid genotype (g), and let (***g*^*^*s_v_***) denote the observed genotype. We defined (***d(g,g’***) ϵ {**0,1,2**}) as the number of unmatched alleles between two unordered diploid genotypes. The expected miscall was

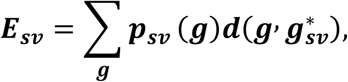

and the hard miscall was

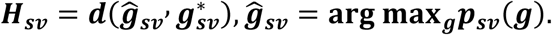

A hard miscall of 0, 1 or 2 therefore indicates that both, one or neither of the predicted alleles matched the observed genotype, respectively.

For each variant, we calculated the mean and standard deviation of both scores across held-out donors, together with the fraction of donors for whom (***H_sv_* > 0**). We also calculated two probability-based measures where indicated: the mean negative log-likelihood of the observed genotype and the mean log-likelihood margin. For each donor, the log-likelihood margin was defined as the log-likelihood of the observed genotype minus that of the most probable alternative genotype.

Mean hard miscall across donors was the primary variant-level measure of residual difficulty and the residual-difficulty feature used in the cis-expression elastic-net models. Donor-level hard miscalls were used to calculate population-level error rates. Mean expected miscall across donors was evaluated as a sensitivity measure. Concordance between mean hard and expected miscall was assessed using Spearman correlation, and cis-eQTL enrichment under the expected-miscall measure was evaluated using a one-sided Mann-Whitney U test.

For each variant, we additionally recorded the number of contributing donors, the number of donors in each genotype class, the MAF estimated from these genotypes and the distance to the nearest TSS annotated in GENCODE v39.

### Variant matching for functional enrichment

To test whether residual difficulty was elevated at regulatory variants while controlling for frequency and genomic location, we used molecular-QTL benchmark datasets released with Borzoi. For cis-eQTL analyses, positive variants were defined by fine-mapped PIP (≥ **0.9**). The corresponding Borzoi negative set served as the candidate control pool.

From this candidate control pool, we performed additional matching separately for Mendel and Evo 2 after restricting both the positive and negative sets to variants with residual-difficulty scores available for the model being analyzed. Variants lacking a MAF estimate or TSS distance were excluded. Matching used two features: held-out MAF and

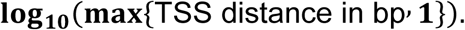

Both features were standardized using the combined positive and candidate-control set for the corresponding model.

Positive variants were processed sequentially and matched greedily, without replacement, to the nearest candidate control in the standardized two-dimensional feature space. A match was retained only if the absolute difference in MAF was no greater than 0.05. If the nearest candidate had already been used or failed this caliper, the positive variant was excluded rather than being matched to the next-nearest candidate. This procedure produced 932 matched positive-control pairs for Mendel and 1,210 pairs for Evo 2. In the Mendel set, mean MAF was 0.2170 in both groups, and mean distance to the nearest TSS was 11,814 bp among positive variants and 11,787 bp among controls. The separately generated random-background variants were retained without matching and were used only as an unpaired descriptive reference.

Splicing-QTLs (sQTLs), polyadenylation-QTLs (paQTLs), and intronic-polyadenylation-QTLs (ipaQTLs) analyses used the positive and negative sets distributed with the Borzoi benchmark. For each positive variant, candidate negatives were identified through the benchmark pairing field linking the negative record to the corresponding positive-variant identifier. One available negative was selected for each positive using random seed 42. After intersection with variants having Mendel residual scores, the analyses included 662 splicing-QTL pairs, 769 polyadenylation-QTL pairs, and 788 intronic-polyadenylation-QTL pairs. Because tissue membership was unavailable for the splicing-QTL subset, these variants were analyzed as a pooled set. Unlike the cis-eQTL analysis, the sQTL, paQTL, and ipaQTL analyses retained the positive-negative pairs defined in the Borzoi benchmark and did not rematch them on MAF or distance to the nearest TSS. The released controls accounted for gene expression and the relevant mechanism-specific site distance.

### Variant prioritization

Separate random forest classifiers were trained using features derived from Mendel and Evo 2 to distinguish cis-eQTLs from their matched benchmark controls. Each classifier comprised 200 trees and used a maximum tree depth of 12, a minimum of five variants per terminal leaf, and a random seed of 42.

For each variant, the residual feature set comprised 11 summaries calculated across donors. Expected miscall, negative log-likelihood, and the Δlog-likelihood margin were each summarized by the mean, median, and inter-donor standard deviation (nine features). Hard miscall was summarized by its mean across donors and the proportion of donors with a value greater than zero (two features). The covariate set comprised MAF, untransformed distance to the nearest TSS, and chromosome indicators. For each set of Mendel- or Evo 2-derived residuals, we evaluated three classifier specifications: residual features only, covariates only, and residual features combined with covariates. Before cross-validation, missing values were imputed using the median of the corresponding feature across the matched analysis set.

Classifier performance was evaluated using stratified fivefold cross-validation. Folds were shuffled using a random seed of 42. Each variant was assigned an out-of-fold predicted probability by the classifier trained on the other four folds. Labels and out-of-fold probabilities were then pooled across folds to calculate the AUROC and AP.

### Cohort-size dependence

To assess how the number of donors contributing to the residual estimates affected variant prioritization, we repeated the residual-only analysis after sampling (***n*** ∊ {**8,17,26,34,42,51,59,68,76,85**}) of the 85 held-out donors without replacement. For each run, the residual features defined above were recomputed from the randomly selected donor pool.

Unlike the primary fivefold evaluation, this sensitivity analysis used a fixed training-test partition. Separately for Mendel and Evo 2, the corresponding matched variant set was divided once into class-stratified training (80%) and test (20%) partitions under seed 42. This partition was held constant across cohort sizes and replicates. Missing values were imputed using medians estimated from the training partition, after which the residual-only random forest described above was fitted and evaluated on the fixed test variants.

Each cohort size was evaluated in five replicate runs, with deterministic, replicate-specific seeds controlling both donor sampling and random-forest fitting. AUROC and AP were summarized by their standard deviation across runs. At n = 85, every run included the complete donor cohort; variation at this point therefore reflects random-forest stochasticity rather than donor subsampling.

### Correlation with regulatory effect metrics

Variants were annotated with cis-eQTL effect estimates from the GTEx v10 significant variant-gene association files. When a variant was significant in more than one gene-tissue association, we retained the normalized effect-size estimate with the largest absolute magnitude. Fine-mapping evidence was obtained separately from GTEx Sum of Single Effects (SuSiE) results by retaining the maximum PIP observed for each variant across all available gene-tissue associations. Consequently, the selected effect-size estimate and maximum PIP were variant-level summaries and were not required to originate from the same gene-tissue association.

Associations between Mendel residual difficulty and regulatory-effect magnitude were evaluated among the model-specific matched cis-eQTL positive variants used in the enrichment analysis. Associations with fine-mapping confidence were evaluated separately among all evaluated Mendel-positive cis-eQTL variants before control matching. Each analysis used its own complete-case set. Spearman rank correlation was used to quantify the association between residual difficulty and either the absolute normalized effect size or maximum PIP, with two-sided (P) values reported.

### Penalty-weighted elastic-net models of gene expression

We evaluated whether Mendel residual difficulty improved cis-expression prediction by fitting four elastic-net models for each evaluable gene-tissue pair: an unweighted baseline, a Mendel-weighted model, a CADD-Phred-weighted model and a model combining the Mendel and CADD-Phred priors. All four models used the same genotype-dosage design matrix, with genotypes encoded as 0, 1 or 2 copies of the alternative allele. The models differed only in the penalty factors applied to individual variants. Expression phenotypes were obtained from precomputed tissue-specific GTEx v10 residual-expression matrices.

Candidate genes and variants were obtained from the frozen cis-variant map used in the analysis. The map-construction pipeline selected genes represented in the expression input, excluded pseudogenes and assigned biallelic SNVs to transcription-start-site-centred cis intervals. The resulting map contained 110 candidate genes. Before model fitting, genomic positions associated with more than one distinct variant identifier were excluded.

Mendel residual difficulty was represented by the donor-level hard-miscall count for each variant, defined as the number of incorrectly predicted alleles and taking values of 0, 1 or 2. CADD-Phred annotations were obtained through FAVOR. Non-numeric, missing and unmatched CADD-Phred values were replaced with the median of the available numeric annotations before merging with the donor-level data. Thus, the model labelled “FAVOR” used CADD-Phred as its annotation-derived prior.

Donor eligibility was determined separately for each gene-tissue pair based on available aligned expression and genomic data; the analysis was not restricted to the 85-donor test cohort. For each gene-tissue pair, expression, genotype dosage, Mendel residual difficulty and CADD-Phred data were aligned by donor. Genes were required to have at least 10 expression records and at least 20 donors shared across the aligned inputs. Before cross-validation, variants were retained when genotype dosage, Mendel residual difficulty and CADD-Phred data were available for at least 80% of aligned donors. At least two variants were required for model fitting. After filtering, 105 genes contributed 4,896 evaluable gene-tissue pairs across 50 tissues.

Penalty factors were constructed separately within each outer training fold. For the Mendel-weighted model, the prior score for a variant was its mean hard-miscall count across donors in that outer training fold. For the CADD-Phred-weighted model, the prior score was the variant’s CADD-Phred annotation. Within each gene, prior scores were converted to empirical cumulative ranks. Ranks were bounded below at 0.05, inverse-squared and normalized so that the mean penalty factor was one:

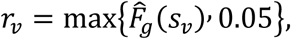

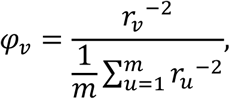

where *(s_v_)* is the prior score for variant (v), (*F^^^*_*g*_) is the empirical cumulative distribution of prior scores among the (m) variants retained for gene (g), and (*φ_v_*) is the resulting penalty factor. Higher prior scores therefore produced smaller penalty factors and less shrinkage. The baseline assigned every variant a penalty factor of one. For the combined model, the separately normalized Mendel and CADD-Phred penalty vectors were averaged with equal weight and renormalized to a mean of one.

Models were evaluated using five repeats of nested cross-validation. Each repeat comprised five outer folds for performance evaluation and five inner folds for selection of the regularization parameter (*λ*). Outer- and inner-fold assignments were shared across the four strategies within each gene-tissue pair and repeat.

Within each outer fold, missing genotype dosages were replaced with variant means estimated from the outer training donors, and the same means were applied to the held-out donors. Missing prior values were similarly replaced using means estimated from the outer training data. Variants with no genotype variation in the outer training set were removed from all four strategies for that fold. Genotype predictors were standardized during model fitting. The elastic-net mixing parameter was fixed at (α = 0.5), and (λ_min_), corresponding to the minimum inner-cross-validation error, was selected separately for each strategy. The initial 80% data-availability filter was applied before the outer cross-validation split. The analysis used a global random seed of 1.

For each repeat, predictions from the five held-out outer folds were concatenated and compared with observed expression using Pearson correlation, producing a repeat-specific cross-validated correlation *(r_j_*). Performance was summarized as

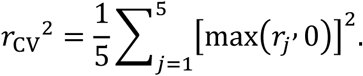

This quantity is the mean squared, non-negative cross-validated Pearson correlation across the five repeats. It is distinct from the predictive coefficient of determination (1 - SSE/SST), which was also calculated but was not used for this performance summary.

For significance assessment, the two-sided Pearson-correlation (P) values from the five repeats were summarized by their median. Within each tissue and modelling strategy, these median (P) values were adjusted across genes using the Benjamini-Hochberg procedure.

The baseline-Mendel comparison used the union of genes with *(q* ≤ 0.05) under either the baseline or Mendel-weighted model and summarized the mean (*r_CV_^2^)* for each model within this paired set. We also calculated the model-specific numbers of genes meeting the FDR threshold. The Mendel-Cadd-Phred comparison used the corresponding FDR union for the Mendel- and CADD-Phred-weighted models.

Tissue selection for the gene-level comparison began with the baseline-Mendel FDR union. Within this set, a gene was eligible when its mean cross-validated Pearson correlation across repeats exceeded 0.10 under either model, and tissues with at least 30 eligible genes were considered. Tissues were ranked first by the excess number of genes passing this threshold under Mendel relative to the baseline, then by the median Mendel-baseline difference in (r_CV_2) across the tissue’s FDR-union set and finally by the number of eligible genes. This procedure selected Colon Transverse.

### Statistical analysis

Bootstrap confidence intervals were calculated using the percentile method. The 95% confidence interval for micro-averaged allele-prediction accuracy was estimated from 1,000 bootstrap resamples of donors. Each sampled donor contributed all scored events; when a donor was sampled more than once, those events were included once per draw. MAF- and genotype-stratified confidence intervals were also based on 1,000 donor resamples; donor-specific accuracies within each stratum were resampled and averaged.

For the random forest classifiers, 95% confidence intervals for the AUROC and AP were calculated from 2,000 bootstrap resamples of the variant labels and their corresponding pooled out-of-fold prediction probabilities. Classifiers were not refitted within bootstrap replicates.

Overall comparisons among cis-eQTL variants, matched controls, and background variants used two-sided Mann-Whitney U tests. MAF- and TSS-distance-stratified comparisons used one-sided tests under the alternative that residual difficulty was greater for cis-eQTL variants than for controls. Spearman correlations were accompanied by two-sided asymptotic P values.

Per-donor accuracies for Mendel and Evo 2 were compared using a one-sided Wilcoxon signed-rank test under the alternative that Mendel was more accurate. A paired, one-sided Wilcoxon signed-rank test was used to assess whether gene-level ΔCV r^2^ was greater than zero. Tissue-level gains, counts of FDR-significant genes, and paired tissue-level model comparisons were reported descriptively; no inferential P values were displayed for these analyses.

Accuracy resampling used a random seed of 42. Variant-level tests, random cross-validation folds and bootstrap confidence intervals did not account for correlations among variants arising from LD.

## Data availability

GTEx v8 whole-genome sequences are available through Database of Genotypes and Phenotypes (dbGaP) under accession [phs000424]. GTEx cis-eQTL summary statistics and fine-mapping posteriors are available through the GTEx Portal. FAVOR annotations were obtained from their respective public distributions at the access dates listed in Supplementary Table S3.

## Code availability

Analysis code and frozen configurations supporting the reported results are available from the corresponding authors upon reasonable request.

---

**Extended Data Figure 1 | Macro-averaged allele-prediction accuracy.** Variant-level macro-averaged allele-prediction accuracy for Evo 2 and Mendel across eligible held-out variant events, giving each variant equal weight.

**Extended Data Figure 2** | Allele-overlap score stratified by diploid genotype class and MAF. Allele-prediction accuracy on held-out individuals partitioned by true genotype class (homozygous minor, heterozygous or homozygous major) within rare (0-1%), low-frequency (1-5%) and common (≥ 5%) MAF bins, shown for (a) Evo 2 and (b) Mendel. Mendel homozygous-genotype accuracy is near ceiling across the MAF spectrum, whereas heterozygote allele-prediction accuracy rises from approximately 0.50 in the rare bin to 0.92 in the common bin. Evo 2 remains near its overall accuracy across genotype classes and MAF bins.

**Extended Data Figure 3 | Memorization control and cohort-size sensitivity.** (a) Allele-prediction accuracy of a descriptive homozygous-major-allele predictor, stratified by MAF bin and true diploid genotype class. The major allele was estimated from deduplicated held-out truth records; this is not a prospective predictor. (b) AUROC and AP for cis-eQTL prioritization as a function of the number of held-out individuals contributing to residual-difficulty estimates. A fixed stratified variant train-test split was used at every cohort size, and error bars show standard deviations across five donor-subsampling replicates.

**Extended Data Figure 4 | Full residual-difficulty distribution and variant-level prioritization summaries.** (a) Full distribution of Mendel mean hard miscall for cis-eQTLs, MAF- and TSS-distance-matched low-PIP controls and random background variants. (b) Per-chromosome AUROC obtained by restricting pooled out-of-fold predictions to each chromosome, comparing Mendel and Evo 2 using residual features alone or combined with MAF, distance to the nearest TSS and chromosome indicators. These are within-chromosome summaries of shuffled variant-level cross-validation and are not chromosome-held-out estimates. (c) Pooled Evo 2 ROC and precision-recall curves comparing residual features alone with residual features plus covariates.

**Extended Data Figure 5 | Covariate balance after cis-eQTL matching.** Distributions of (a) minor-allele frequency and (b) log-transformed distance to the nearest TSS for high-PIP cis-eQTLs, matched low-PIP controls and the unpaired random-background reference in the Mendel analysis.

**Extended Data Figure 6 | Residual-difficulty enrichment extends to sQTL, paQTL and ipaQTL variants.** (a-c) Per-variant mean hard miscall under Mendel for QTL positives, exact released Borzoi-matched controls and an unpaired random background reference, shown for (a) sQTLs, (b) paQTLs and (c) ipaQTLs. Complete released pairs were retained when negative INFO/PI equalled positive ID, with one locally scored negative selected per positive using seed 42; the analysis included 662 pooled sQTL pairs, 769 paQTL pairs and 788 ipaQTL pairs. The sQTL analysis is pooled because tissue-specific pair membership was not retained in the merged release. Released controls account for gene expression and mechanism-specific site distance; no additional MAF or TSS-distance matching was applied. (d-f) Proportions of variants in each mean-hard-miscall category for the corresponding sQTL, paQTL and ipaQTL sets.

**Extended Data Figure 7 | Associations of residual difficulty with allele frequency and genomic context.** Mendel mean hard miscall as a function of (a) MAF, (b) log-transformed distance to the nearest TSS and (c) local guanine-cytosine content within ±200 bp. Points show binned means, shaded regions show approximate 95% confidence intervals and labels report Spearman ρ calculated across individual variants.

**Extended Data Figure 8 | Expected-miscall sensitivity analysis.** (a) Concordance between mean expected miscall and mean hard miscall across Mendel-scored variants. Color indicates the number of variants per hexagonal bin, and the dashed line denotes equality. (b) Expected miscall for cis-eQTLs, MAF- and TSS-distance-matched low-PIP controls and an unpaired random-background reference. Boxes show interquartile ranges, whiskers show the 5th-95th percentiles and diamonds denote means; the displayed P value is from a one-sided Mann-Whitney U test comparing cis-eQTLs with controls.

**Extended Data Figure 9 | Training of chromosome-specific Mendel models.** Overview of the per-chromosome training workflow, optimization schedule, validation-loss monitoring and checkpoint-selection procedure.

## Notes

### Competing Interest Statement

The authors have declared no competing interest.

