## Supplementary figures and images for "Mendel, a foundation model of human genetic variation, prioritizes regulatory variants and improves gene expression prediction"

### Extended Data Figure 1

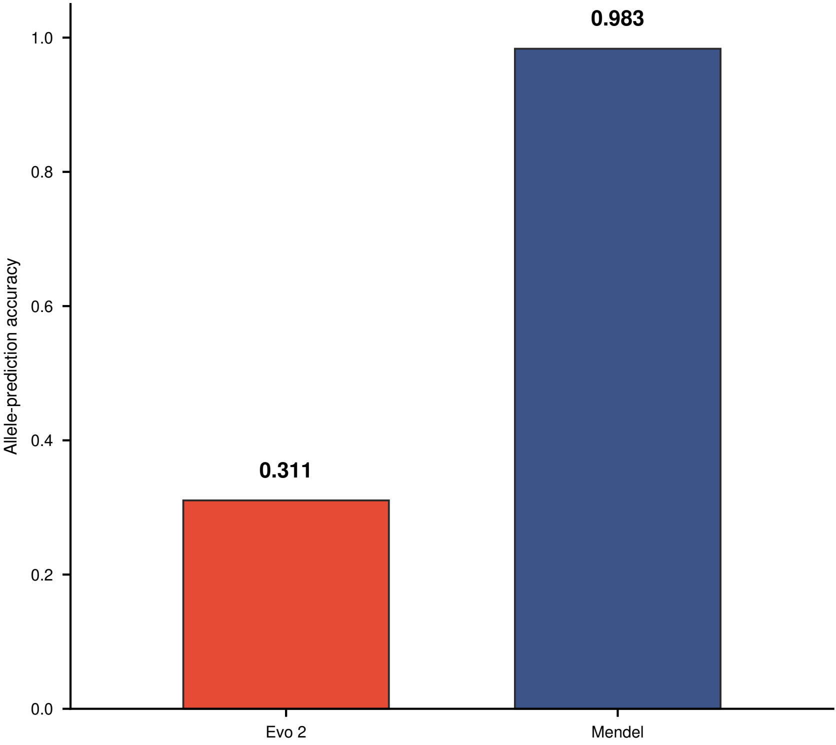

### Extended Data Figure 2

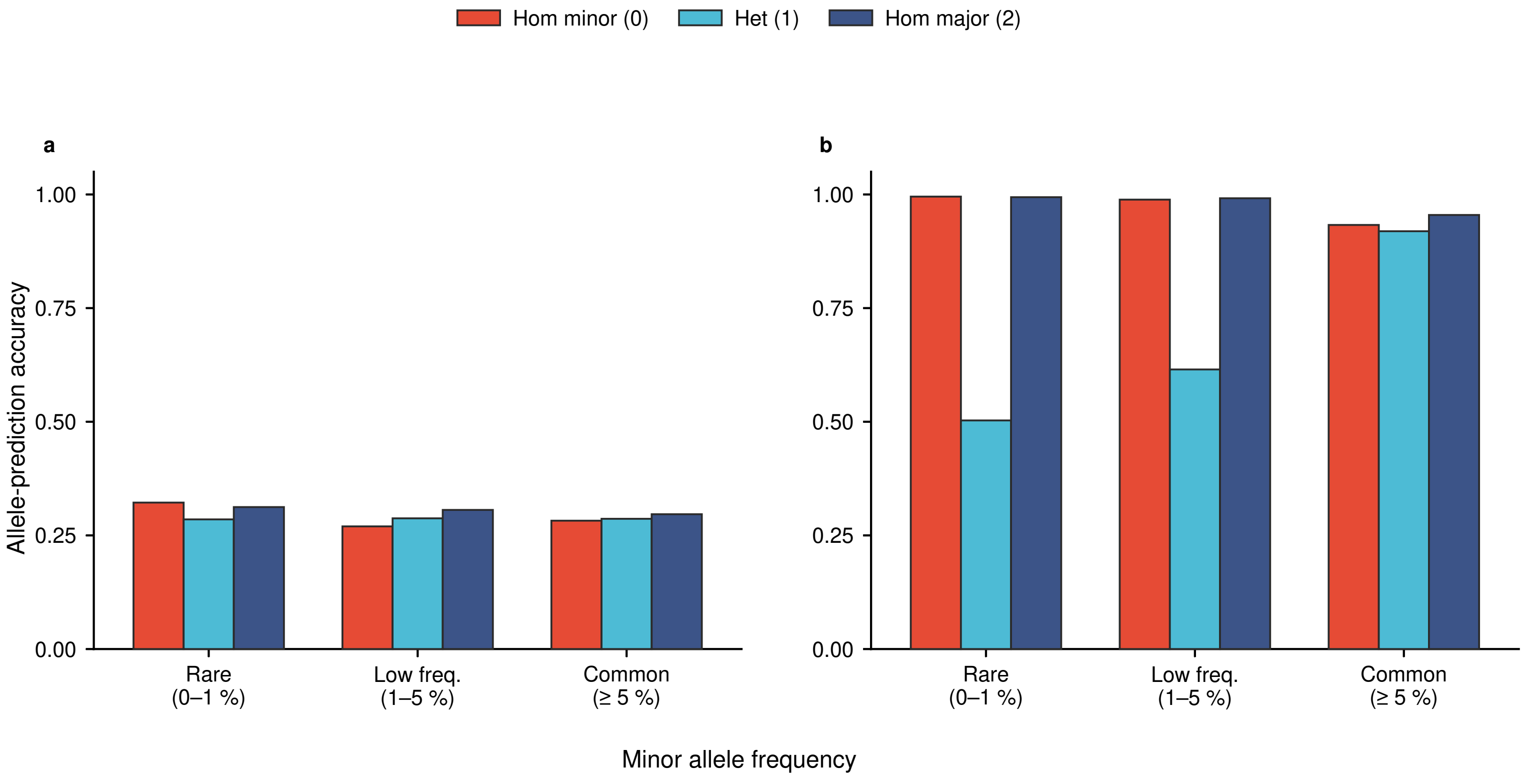

### Extended Data Figure 3

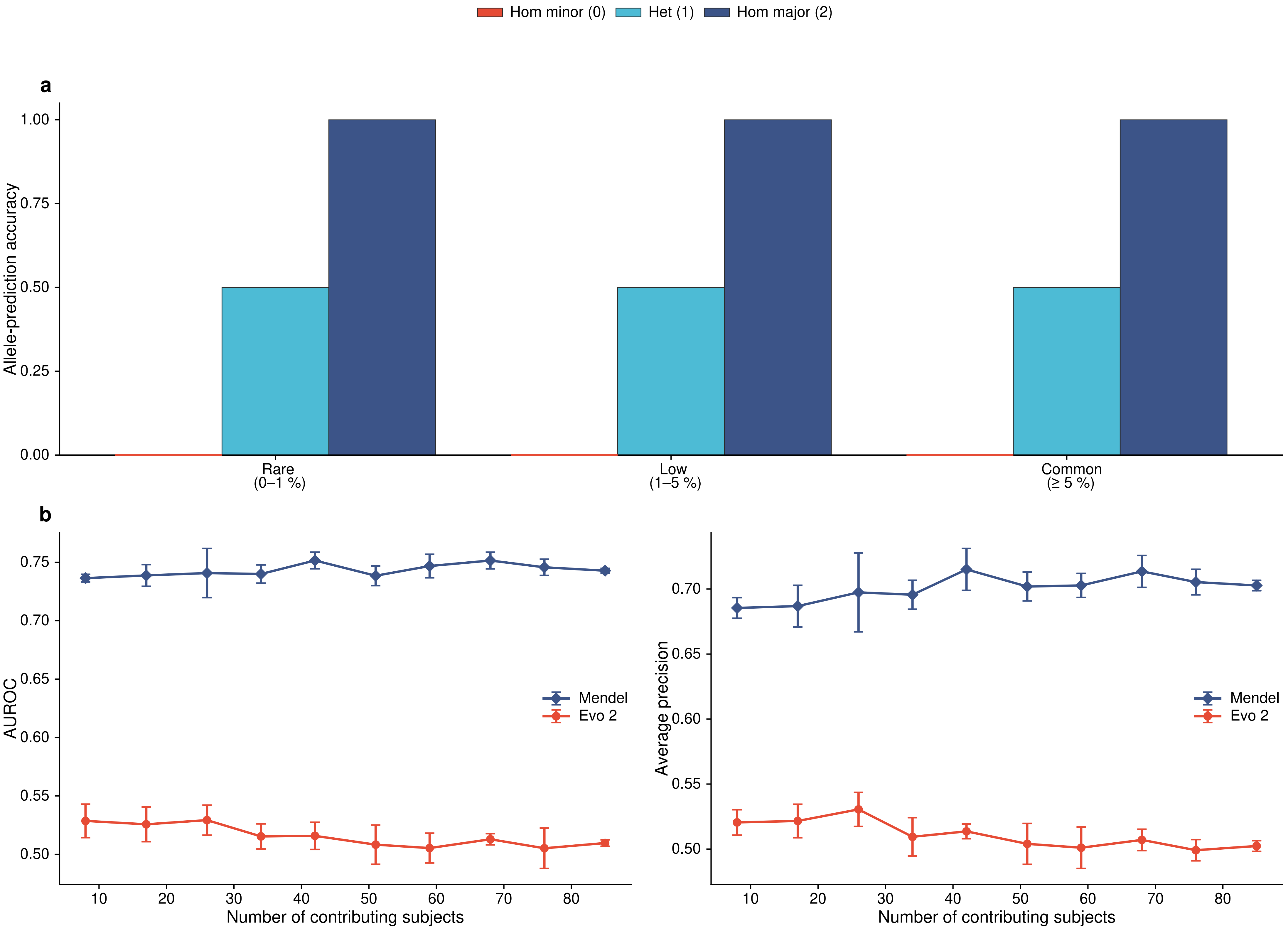

### Extended Data Figure 4

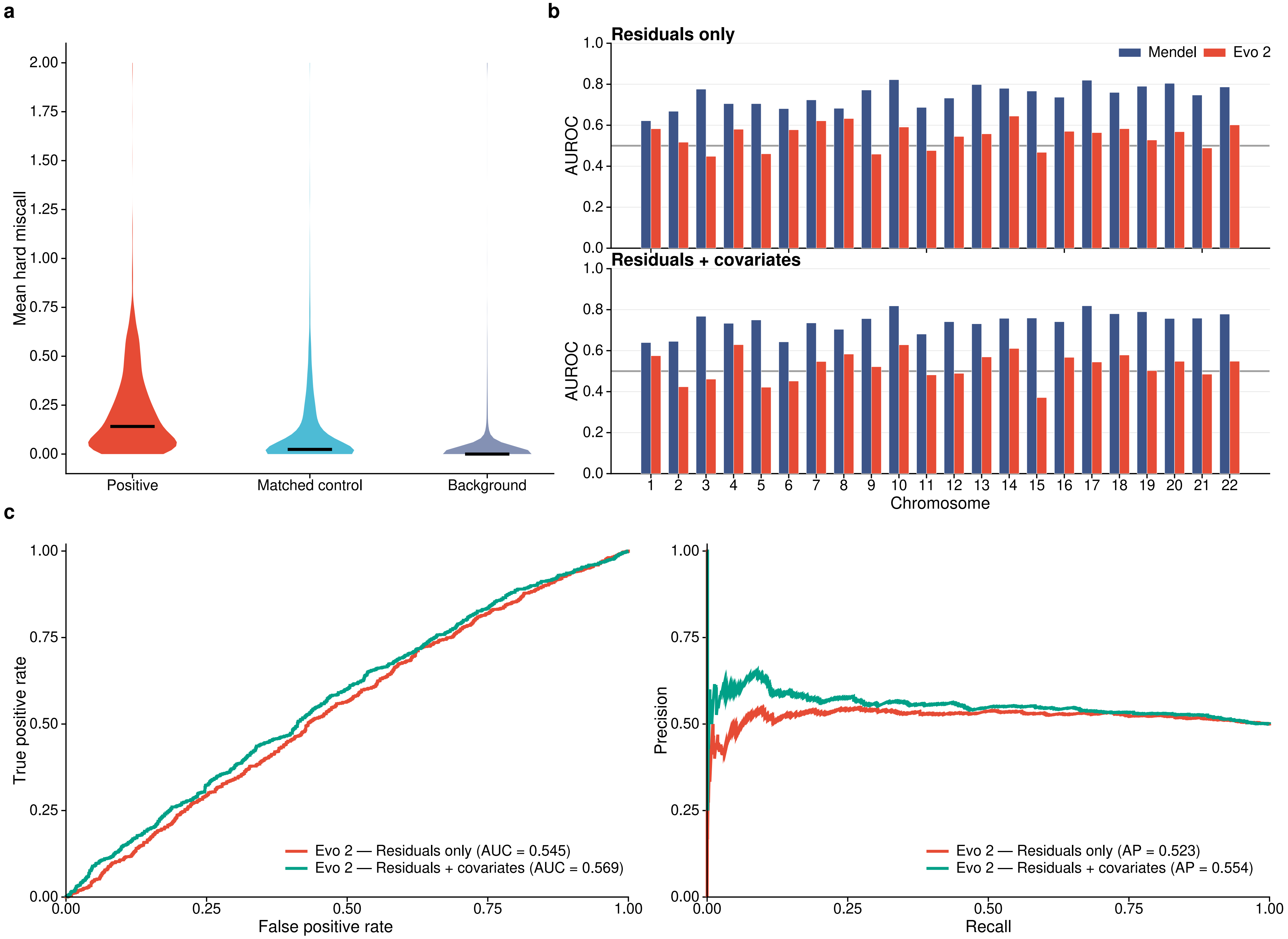

### Extended Data Figure 5

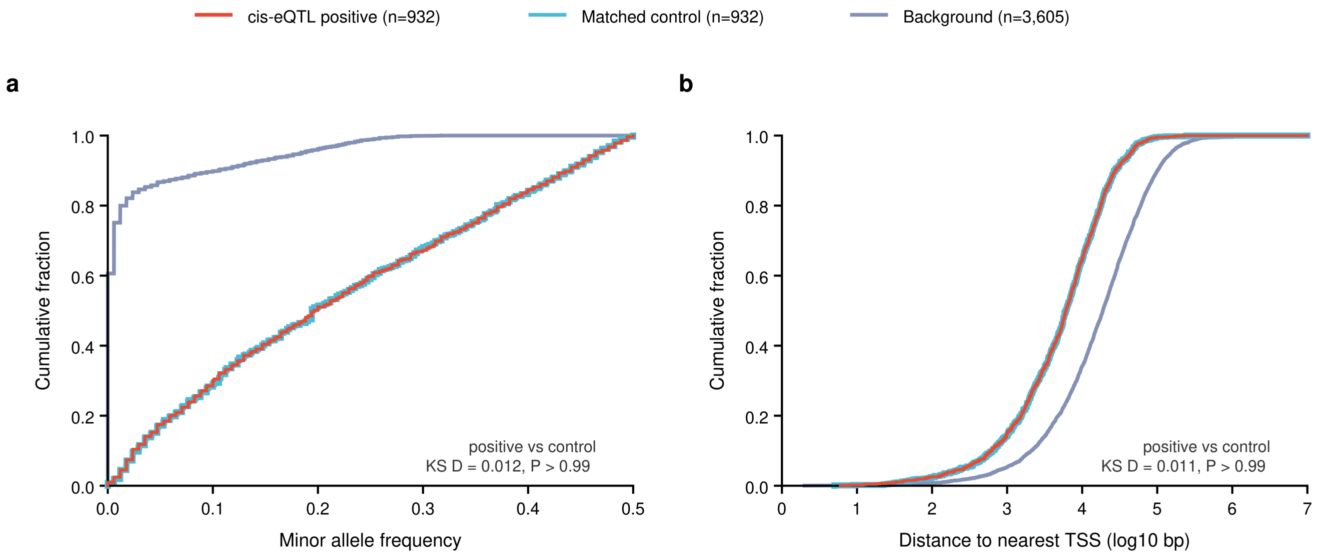

### Extended Data Figure 6

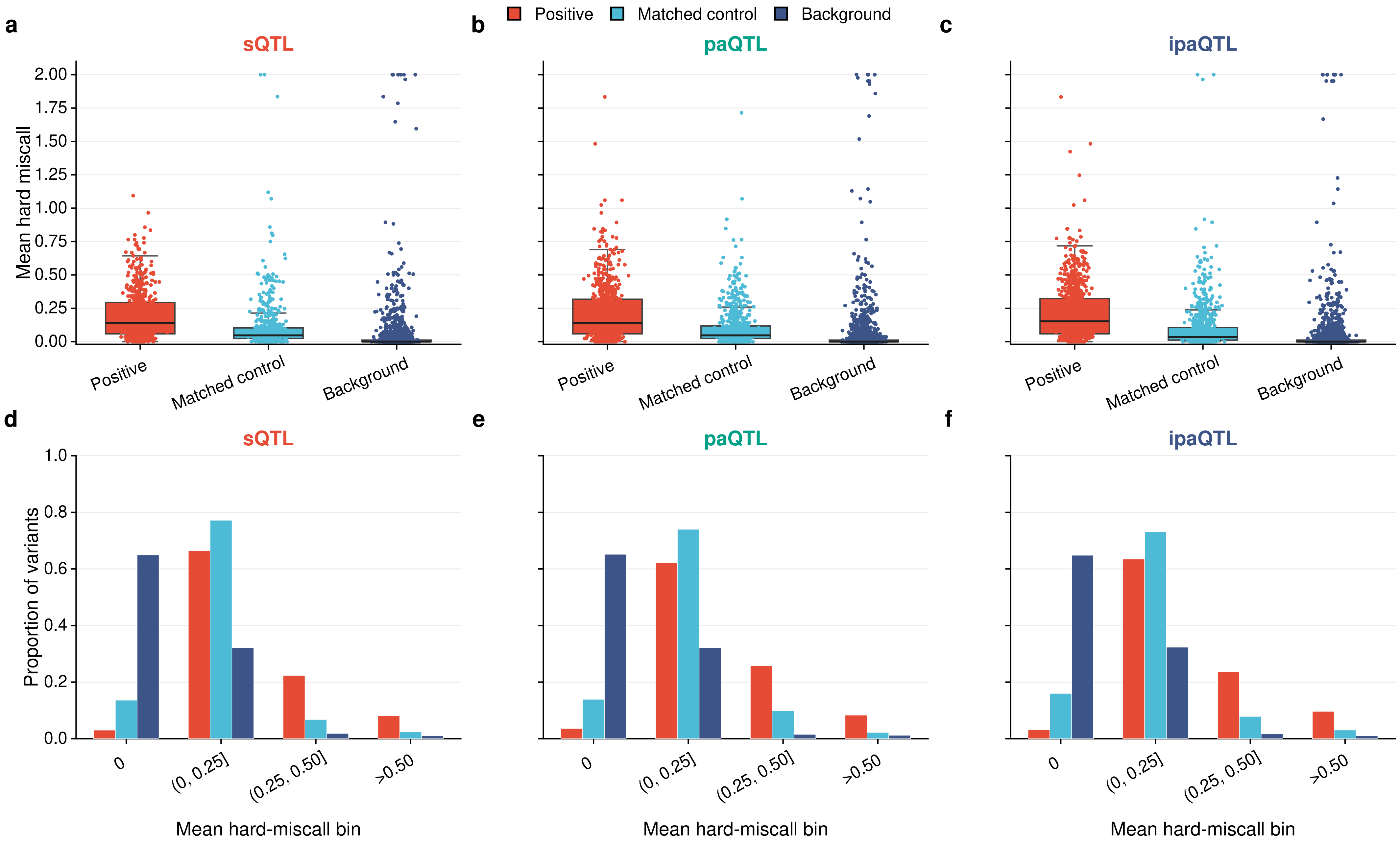

### Extended Data Figure 7

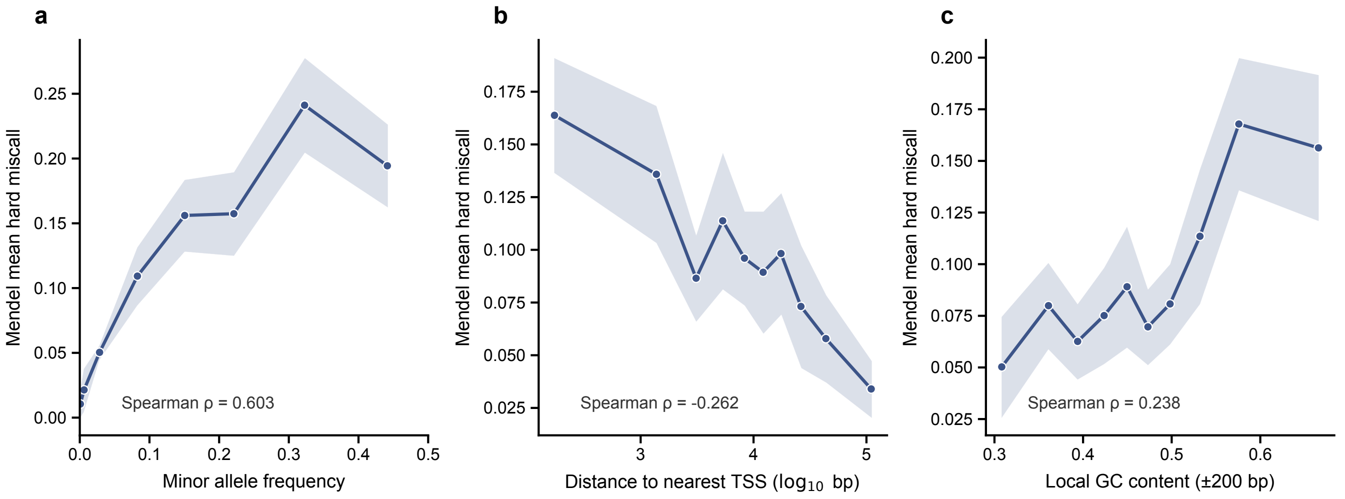

### Extended Data Figure 8

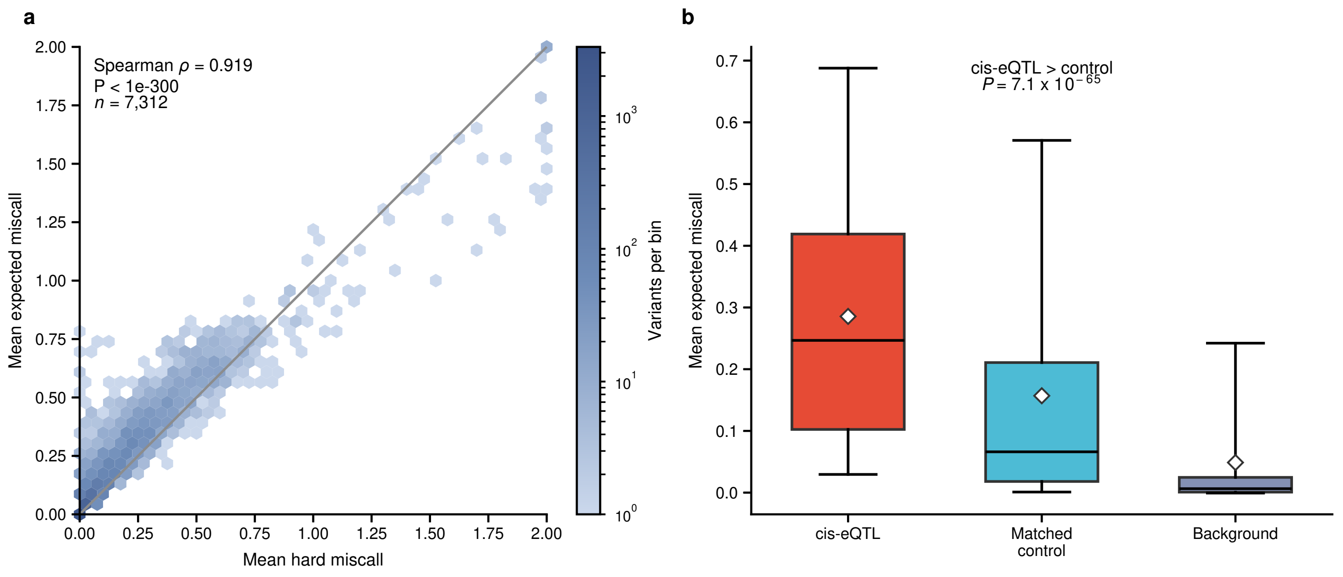
