## Extended Data Figure 9 for "Mendel, a foundation model of human genetic variation, prioritizes regulatory variants and improves gene expression prediction"

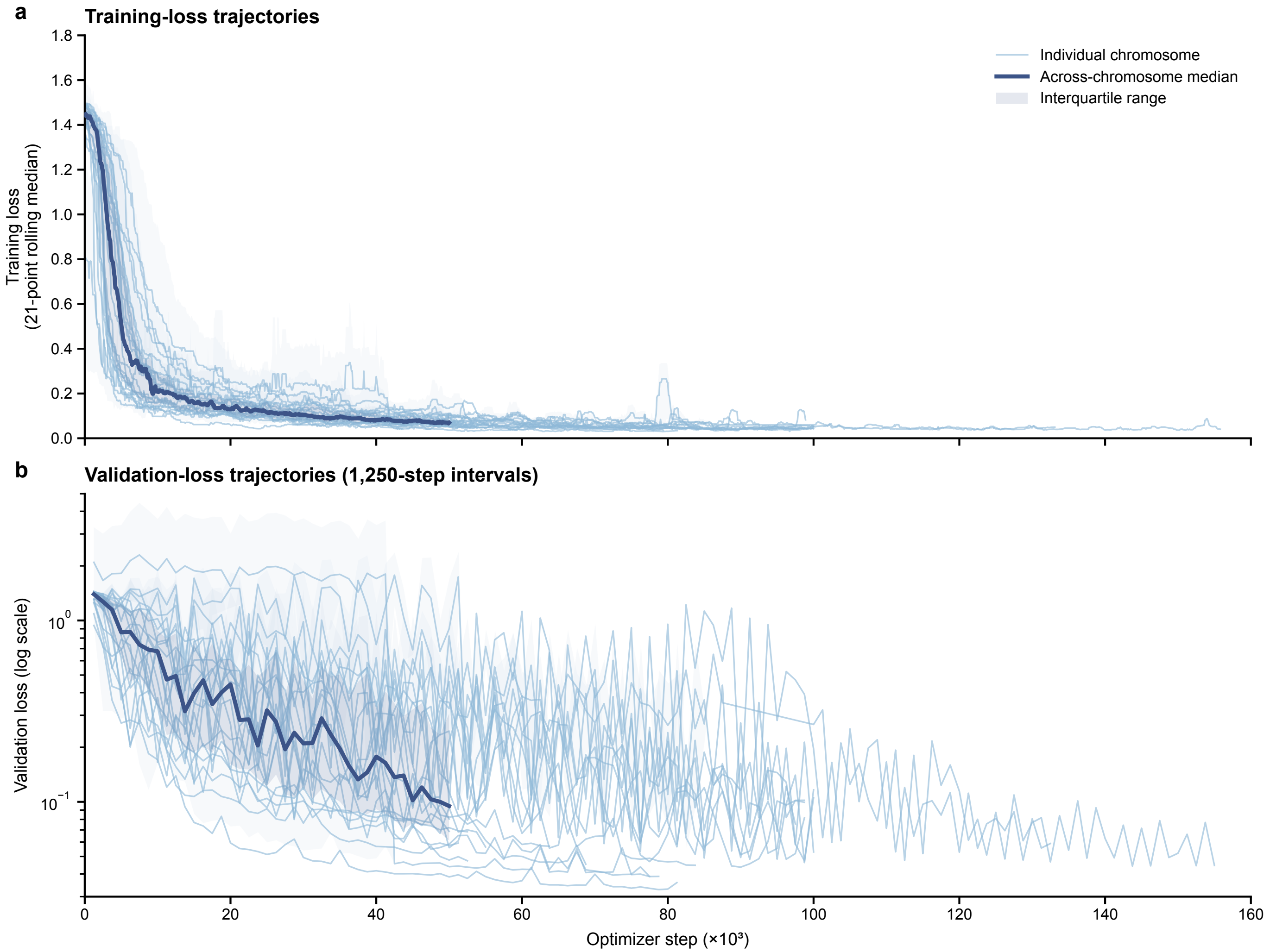


Median and interquartile range are shown through 50,000 optimizer steps, where all 22 chromosome models contribute.
W&B exports dated 20 August 2026; faint ribbons show min–max ranges for grouped chr10, chr11, chr12 and chr22 traces, and one chr14 validation point.
